# Detection of CRISPR-Cas-induced mutations in *Daphnia*

**DOI:** 10.64898/2025.12.05.692657

**Authors:** Swatantra Neupane, Michael E. Pfrender, Li Wang, Sen Xu

## Abstract

CRISPR-Cas9 has established itself as a robust tool for conducting loss of function gene research in emerging model species including the freshwater zooplankton *Daphnia*. However, sensitive detection of mutations, especially in genetic mosaic and pooled samples, remains a challenge. In this study we evaluate two of the most widely used mutation screening techniques, the T7 Endonuclease I (T7EI) assay and Fragment Analysis (FA) for their sensitivity, accuracy, and practical use in detecting CRISPR-induced indels in four targeted genes, *DNMT3A*, *DNMT3B*, *PERIOD2*, and *DMRT1* in *Daphnia magna*. Here, we show that T7EI, although it offers a quick and cost-effective screening method, often produces false positives, especially when examining pooled samples. Conversely, FA facilitates detecting allele size differences at a fine resolution, reproducibility in detecting indels, and distinguishing zygosity and is more reliable as a method to detect mutation. Our comparative analyses convey the importance of carefully selecting the appropriate screening methods depending on research questions.

## Introduction

The CRISPR-Cas gene-editing system has become a leading method for generating targeted genetic modifications in various animal models (Hsu et al. 2014; Lee et al. 2020). Cas-induced DNA double-stranded breaks (DSBs) are repaired by error-prone mechanisms, e.g., non-homologous end joining (NHEJ), or homologous recombination when a repair template is provided (Aird et al. 2018; Huang and Puchta 2019). While significant advancements have been made in optimizing in-vivo delivery techniques and editing efficiencies (Luther et al. 2018; Demirci et al. 2022), efficient and reliable identification of CRISPR-edited organisms, or crispants, remains a time consuming and tedious task.

Distinguishing between crispants and unedited individuals within a mixed population requires robust mutation detection methods. The most robust and widely used mutant screening methods are next-generation sequencing (NGS) whole-genome sequencing and amplicon sequencing, along with Sanger sequencing. While genomic sequencing is highly accurate and offers a more comprehensive evaluation of genome-wide on-target and off-target induced mutations, scaling it for high-throughput mutant screening is often impractical due to costs and complexity (Vonk et al. 2025). Single-molecule real-time sequencing (SMRT) and nanopore sequencing are emerging as promising alternatives to conventional NGS due to their ability to detect structural variations, long-range haplotypes, and epigenetic modifications in CRISPR-edited genomes (Ardui et al. 2018). Nonetheless, their current costs do not allow them to be applied for large-scale screening.

Alternatively, fluorescence-based Assays (FA), which involves the use of fluorescently labeled dyes or proteins, can be applied *in vitro* for mutation detection, e.g., TaqMan probes, melting curve analysis and real-time PCR (Huang et al. 2011). For example, digital PCR (dPCR) has emerged as a highly sensitive and precise alternative to detect rare mutations and low-abundance CRISPR-induced indels. Unlike traditional qPCR, dPCR partitions the sample into thousands of individual reactions, enabling absolute quantification of mutation events without the need for standard curves (Quan et al. 2018). Similarly, high-resolution melting (HRM) analysis emerged as a cost-effective, post-PCR method for detecting small sequence variations, making it a useful complement to fluorescence-based assays (Wittwer et al. 2003). Another emerging method for mutation detection is amplicon-based CRISPR editing analysis using droplet-based microfluidics, which enhances sensitivity and throughput for detecting rare editing events (Zhao et al. 2023). Moreover, Cas9-induced indel detection by barcode sequencing is a novel approach that uses targeted deep sequencing to provide high-resolution profiles of CRISPR-mediated mutations and their distributions (Bell et al. 2014). These advancements contribute to improving the accuracy and scalability of mutant screening, particularly for large-scale or multiplexed CRISPR experiments. However, screening for a simple CRISPR induced mutations without *in vivo* modification, these new methods seem more labor intensive.

Among the most widely used cost-effective approaches for mutant screening are the T7 Endonuclease I (T7EI) assay and Fragment Analysis (FA). The T7EI assay is a mismatch detection method that identifies insertions and deletions (indels) at the target site by taking advantage of the T7’s ability to enzymatically cleaving mismatched DNA heteroduplexes (Mashal et al. 1995; Youil et al. 1995). It is frequently employed due to its simplicity and rapid processing time. However, T7EI’s sensitivity to heteroduplex formation can lead to false positives or inconsistent results (Babon et al. 1995; Mashal et al. 1995; Youil et al. 1995), particularly in organisms with complex genetic backgrounds (Sentmanat et al. 2018). Additionally, the assay struggles with detecting small indels or mutations present at low frequencies (in cell populations), making it less reliable for accurate mutation characterization (Vouillot et al. 2015).

Fragment Analysis (FA) is a precise and quantitative method for detecting Cas-induced mutations using capillary electrophoresis to distinguish DNA fragments based on size variations caused by indels. FA is advantageous for its quantitative nature and ability to detect CRISPR-induced mutations based on allele size variations of target loci, offering a higher resolution approach for identifying indels. The process begins with PCR amplification of the target locus using fluorescently labeled primers. The resulting amplicons are then separated by capillary electrophoresis, which accurately measures fragment sizes.

FA provides higher resolution than agarose-gel-based methods, enabling precise quantification of indel sizes. Unlike T7EI, it does not depend on heteroduplex formation, enhancing its reliability for detecting subtle mutations. Additionally, FA can differentiate between heterozygous and homozygous mutations, offering insights into mutation zygosity. However, the need for specialized equipment, such as a capillary-based genetic analyzer, restricts its use.

In this study, we compare the efficiency, accuracy, and limitations of T7 Endonuclease I assay and Fragment Analysis for identifying CRISPR-induced mutations in the microcrustacean *Daphnia*. By evaluating their performance in detecting crispants, we aim to provide insights into which methods to choose for in-vivo mutant screening, considering factors such as sensitivity, reproducibility, and practicality.

## Materials and Methods

### Daphnia system, target genes for knockout, and screening strategies

*Daphnia magna* has been a model system for genomics, population and evolutionary studies for several decades. With its whole genome sequenced and well annotated (Chaturvedi et al. 2023), it serves as a good system for undertaking CRISPR experiments. *Daphnia magna,* as with other *Daphnia* species, has a cyclically parthenogenetic mode of reproduction where under favorable conditions females reproduce asexually making embryos without mating. When exposed to adverse conditions female *Daphnia* switch to sexual reproduction where they produce recombinant offspring with males in the populations. The parthenogenetic offspring are genetically identical to their mother. Previously, we showed that successfully edited females may produce offspring of variable mutation profile (mosaicism) at the target sites (Xu et al. 2025).

For our gene knockout experiments we used the *D. magna* isolate LRV01, which has a high quality genome assembly (Chaturvedi et al. 2023). Four genes, *DNMT3A*, *DNMT3B*, *PERIOD2*, and *DMRT1* were targeted for CRISPR/Cas9 knockout mutations, see Table 1.

**Table 1.**
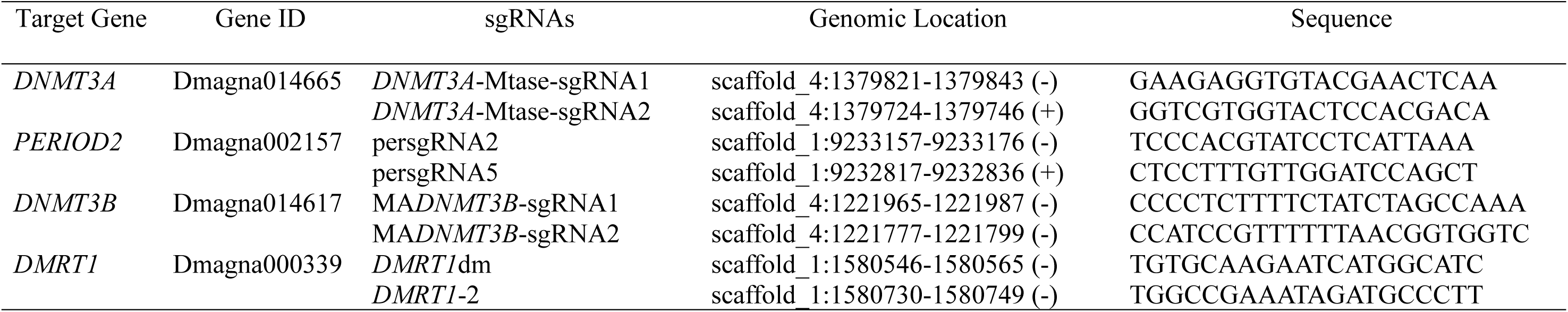
CRISPR target sites and sgRNA target sequences.

**Table 2.**
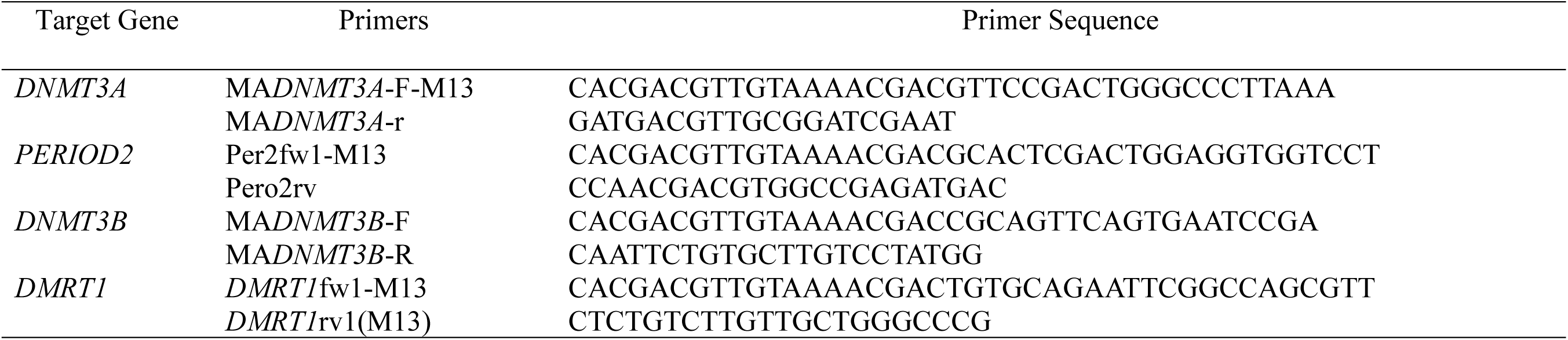
Fragment Analysis PCR primers. The forward primers include M13-tail (5’-CACGACGTTGTAAAACGAC-3’)

We use fragment analysis to resolve subtle differences in target amplicon sizes caused by Cas-induced indels. This method allows us to distinguish between clean, single-signature edits and more complex mixtures which are more prevalent in pooled samples. In parallel, T7 endonuclease assays are performed as a screen for heteroduplex formation, though we’ve observed that T7 can miss certain types of edits or give false positives, particularly in pooled samples. Together, these methods help us piece together the mutation mosaics within Brood 1 and determine whether edits were transmitted through the germline.

### SgRNA design and Cas9 delivery

For each target gene sequence, we used IDT’s custom guide RNA design tool (https://www.idtdna.com/site/order/designtool/index/CRISPR_SEQUENCE) to identify optimal sgRNAs based on their on-target scores, which considers factors such as the surrounding nucleotide sequences, GC composition near the PAM site, and the presence of cytosines within the 20-nucleotide target region. Additionally, sgRNAs were selected based on their genomic location, prioritizing those within exons or protein domains to maximize the likelihood of functional disruptions. A subset of high-scoring crRNAs was then synthesized and assembled into RNP complexes with Cas9 nuclease in vitro. The assembly involved first fusing each crRNA to a tracrRNA (supplied separately) to form a functional tracrRNA-crRNA hybrid (sgRNA), which was subsequently complexed with the Cas9 nuclease to generate a ribonucleoprotein (RNP) complex.

The activity of the assembled RNP was tested in vitro by performing a digestion assay on the PCR amplicon containing the target site. Specifically, a PCR amplicon was designed with the target site positioned off-center, producing two fragments of unequal lengths which upon digestion are easily distinguishable on an Agarose gel. For the digestion assay, 2 μL of PCR amplicon (>150 ng), 1 μL of nuclease reaction buffer, 1 μL of RNP, and 6 μL of water were mixed in a reaction tube. The mixture was then incubated at 37 °C for 1 hour. Then, 1 μL of Proteinase K (20 mg/ml) was added and incubated at 56 °C for 10 minutes to deactivate and decouple the Cas9 enzyme. The digested product was run on a 1 % agarose gel electrophoresis alongside an undigested wild-type (WT) control. Successful RNP cutting activity results in two distinct bands, corresponding to the cleaved fragments, whereas the WT sample shows a single band equal in length to the sum of the digested fragments.

### Microinjection for gene editing

We followed a previously described microinjection procedure for generating mutants (Xu et al. 2025). We selected about 100 female daphniids with dark ovaries, indicating they were close to molting and ovulating. Each was placed in COMBO medium (Kilham et al.1998) with 60 mM sucrose and monitored for molting. We watched for ovulation, which starts 10–15 minutes post-molting. When ∼80% of embryos entered the brood chamber, the female was moved to ice-cold COMBO (1.5 °C) for 6 minutes to slow embryo development. We then dissected the embryos on an inverted Petri dish, aligned them along the edge, and keeping a thin layer of COMBO over the embryos, which are ready for injection.

Injections were performed immediately after embryo preparation. We backloaded 1 μl of RNP into microinjection needles using Eppendorf microloader tips (cat. no. 930001007). Each embryo was injected with 1–2 nl of RNP, released near the center using pressures between 100–220 hPa (background: 100–200 hPa) over 0.8 sec. Post-injection, embryos were incubated in COMBO with 60 mM sucrose for ∼30 min at room temperature, then transferred to 60 mM sucrose COMBO in 96-flat well plates and cultured at room temperature. Hatching typically completes within ∼48 hrs.

### General strategy for screening mutants

Post-hatching, female neonates were cultured in 6-well plates. As these females started to carry embryos, they were individually isolated, with each female placed in a single well in a separate 6-well plate. Offspring from the first brood were distributed into the remaining five wells of the same plate. Once these first brood individuals reproduced (usually took 7-10 days under regular feeding of algae), animals from each of the five wells (each well representing a potentially different genotype) were pooled for DNA extraction and genotyped using fragment analysis or T7 assay for the target genes. Pooled samples showing signs of mutation were further analyzed by genotyping (T7 assay or fragment analysis) individual first brood isolates to identify which of the first brood are mutants.

### T7 Endonuclease I (T7EI) Assay

The T7 Endonuclease I (T7EI) assay is commonly used to detect Cas-induced mutations by leveraging the enzyme’s ability to recognize and cleave mismatched DNA sequences. The target genomic region is first amplified via PCR to produce double-stranded DNA fragments. These amplicons undergo denaturation and reannealing, promoting heteroduplex formation between wild-type and mutant DNA strands when mutations are present. T7EI selectively cleaves these heteroduplexes at mismatch sites, generating distinct DNA fragments that are subsequently analyzed by gel electrophoresis to determine the presence of mutations.

To perform the T7 digestion on pooled as well as individual samples, approximately 200 ng of PCR amplicon was combined with 2 μL of 10X NEBuffer 2 and molecular-grade water to a final volume of 19 μL. Hybridization was achieved by initial denaturation at 95°C for 5 minutes, followed by annealing from 95°C to 85°C at a ramp rate of −2°C/second and from 85°C to 25°C at −0.1°C/second, with a final hold at 4°C. Alternatively, after denaturation at 95°C, the reaction can be allowed to cool gradually at room temperature, eliminating the need for programmed ramp rates. After hybridization, 1 μL of T7 endonuclease (NEB catalog number M0302S, 10 units/µl) was added to the mix, which was then incubated at 37°C for 20 minutes. The reaction was terminated by adding 1.5 μL of 0.25 M EDTA and then analyzed by agarose gel electrophoresis.

### Fragment Analysis (FA)

We employed an M13 fluorescence-tagged PCR (Schuelke 2000) approach for fragment analysis. The forward primer was modified with an M13 tail sequence (CACGACGTTGTAAAACGAC) at its 5′ end. The PCR reaction included the M13-tailed forward primer, a reverse primer, and an M13 sequence labeled with one of the NED, PET, FAM, or VIC fluorescent dyes. The thermal cycling conditions were as follows: initial denaturation at 94°C for 5 minutes, followed by 30 cycles of 94°C for 30 seconds, 60°C for 45 seconds, 72°C for 30 seconds, and a final extension at 72°C for 10 minutes. 2 ul of the PCR product was diluted and used for fragment analysis using LIZ ladder 600 (GeneScan™ 600 LIZ™, Catalog number 4408399) on an ABI 3130 Genetic Analyzer (Life Technologies) at DNA Core Facilities at University of Missouri, Columbia. We used the Osiris software 2.15.1 (NCBI) to score the size of alleles.

## Results and Discussion

Although the T7EI assay is rapid and straightforward, its dependence on heteroduplex formation can result in variable detection efficiency. Small indels, particularly single-nucleotide insertions or deletions, may not create sufficient mismatches for enzyme recognition, leading to false negatives. Additionally, T7EI’s activity is influenced by sequence context, reducing its reliability for detecting certain mutations. These limitations are generally acknowledged by the manufacturer, noting that the enzyme has low sensitivity for mismatches of unknown composition, single base pair mismatches, and may require titration for optimal cleavage. These constraints pose challenges when screening for unknown types of mutations that are potentially induced by Cas enzyme.

### Case Study #1 – *DNMT3A* gene knockout

Nine samples of brood 1 for *DNMT3A* gene knockout were initially screened for Cas-induced mutations using the T7EI assay. Out of the nine, four samples tested positive (Table 3), showing additional bands indicative of potential mutations. To verify these results, whole genome sequencing instead of FA analysis was subsequently performed on the four T7-positive samples. Interestingly, WGS did not detect any mutations in these samples (Table 3). This outcome suggests the occurrence of false positives in the T7EI assay, highlighting its limitations in mutation detection accuracy.

**Table 3.** Mutant screening of 3A1-3A9 for *DNMT3A* gene. Mutant screening was done using T7 endonuclease assay and whole genome sequencing. Positive refers to a mutation detected and negative as no mutation.

| *DNMT3A* | WGS | T7EI |
| --- | --- | --- |
| 3A1 | N/A | Negative |
| 3A2 | N/A | Negative |
| 3A3 | N/A | Negative |
| 3A4 | N/A | Negative |
| 3A5 | Negative | Positive |
| 3A6 | Negative | Positive |
| 3A7 | N/A | Negative |
| 3A8 | Negative | Positive |
| 3A9 | Negative | Positive |

### Case Study #2 – PER2 gene knockout

A total of 25 hatched samples were screened for CRISPR-induced mutations in the Per2 (*PERIOD2*) gene using T7EI. Of these, three tested positive, suggesting potential mutations. To validate these results, Fragment Analysis (FA) was performed on the T7EI-positive samples (Table 4). FA confirmed mutations in only 2 samples on *PERIOD2* gene, revealing one false positive from the T7EI assay for *PERIOD2* (Figure 1).

**Figure 1.**
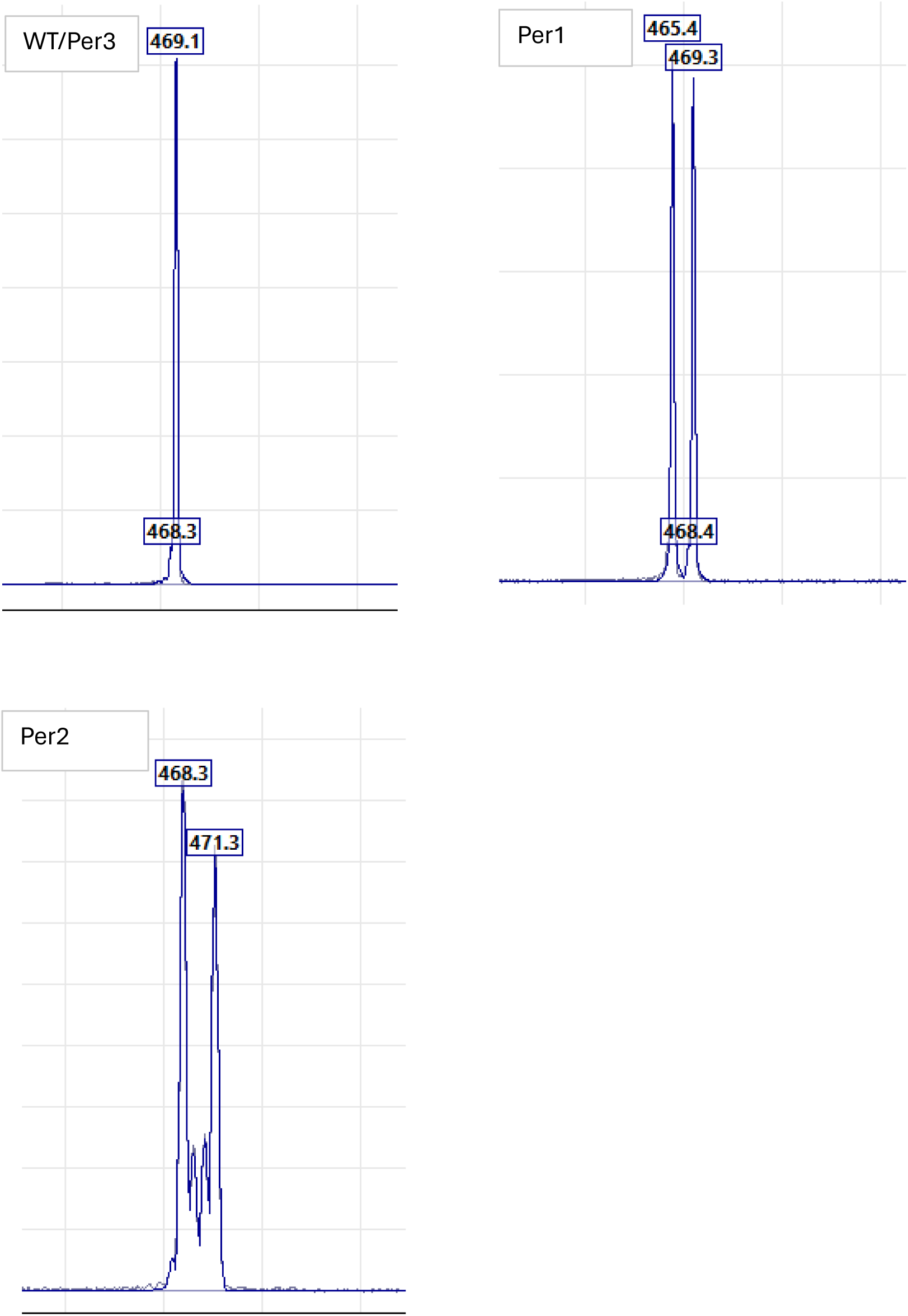
Fragment analysis of Per2 gene, WT allele has a single peak at 469.1, Per1 and Per2 have mutant heterozygous alleles

**Table 4.** Mutant screening of Per-1 to Per-3 for *PERIOD2* gene. Out of 25 samples three tested positive for T7EI assay, two of which were supported by fragment analysis.. Positive refers to a mutation detected and negative as no mutation.

| *PERIOD2* | FA | T7EI |
| --- | --- | --- |
| Per1 | Positive | Positive |
| Per2 | Positive | Positive |
| Per3 | Negative | Positive |

This discrepancy highlights the limitations of T7EI, which can produce false positives due to its reliance on heteroduplex formation. In contrast, FA provided more accurate detection based on amplicon lengths. These findings underscore the importance of validating T7EI results with a more accurate method like FA for reliable mutation screening.

### Case study #3 – *DNMT3B* gene knockout

A total of 9 pooled samples, each representing a brood 1 offspring genotype from knockout experiments targeting the *DNMT3B* gene, were screened for mutations. For each sample, 3–5 brood 1 individuals were pooled for fragment analysis. Out of 9, one sample (S5) showed a distinct PCR amplicon profile (Figure 2) on agarose gel, including a wildtype-like band and an additional higher molecular weight band, suggesting a potential insertion (later verified with fragment analysis). The remaining 8 samples showed wildtype-like bands and were further investigated using both T7 and Endonuclease I (T&EI) and Fragment Analysis (FA). Fragment Analysis revealed mutations in 6 samples: S2, S3, S5, S6, S7, and S9 (Table 5). T7EI cleavage on the same set also detected mutations in these same 6 samples (Figure 3). In this case, T7EI and FA consistently identified the same set of positive samples, supporting their agreement in detecting CRISPR-induced mutations.

**Figure 2.**
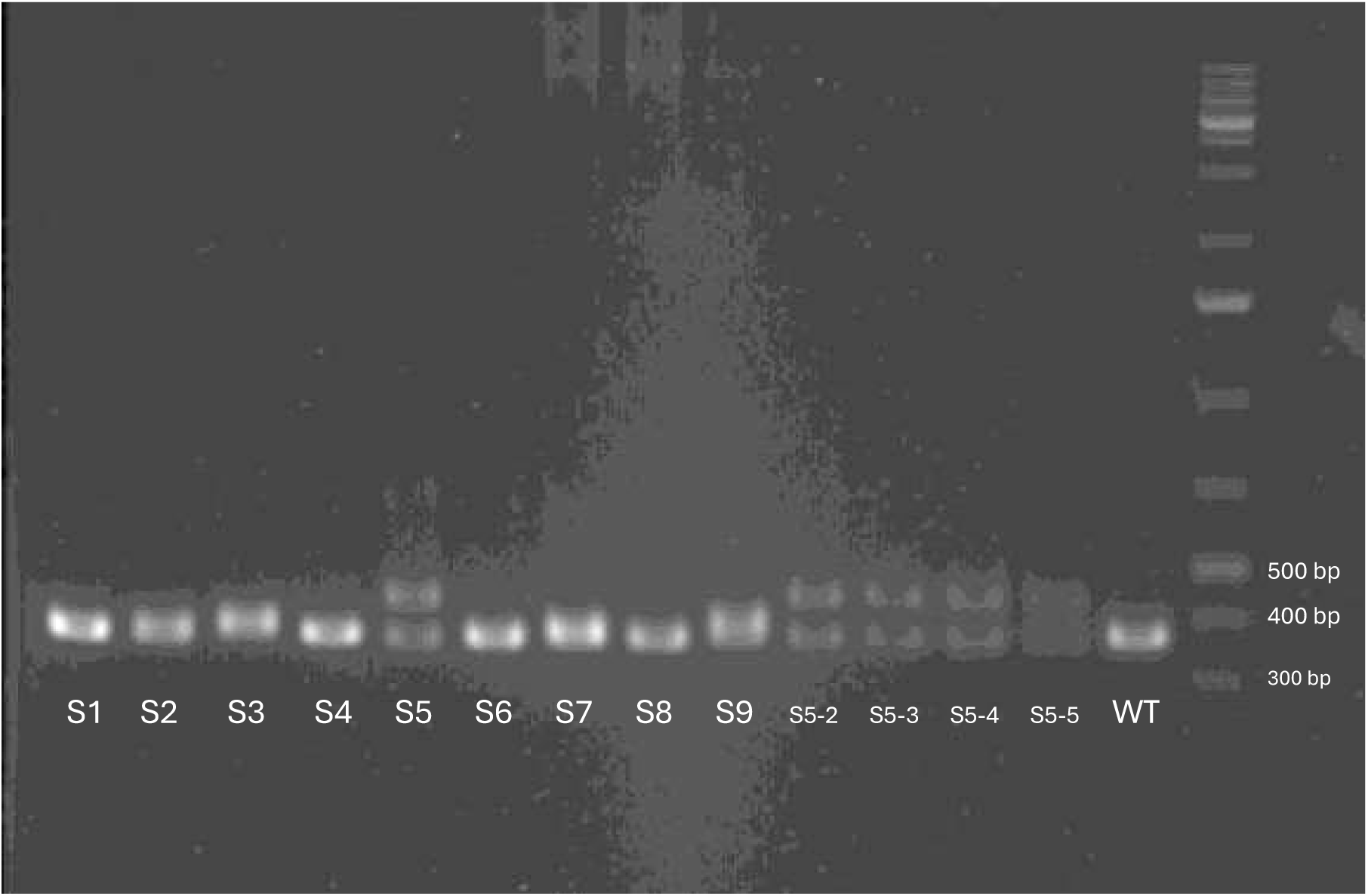
PCR amplification for *DNMT3B* gene of S1-S9, and individual brood 1 genotypes from S5 mom.

**Figure 3.**
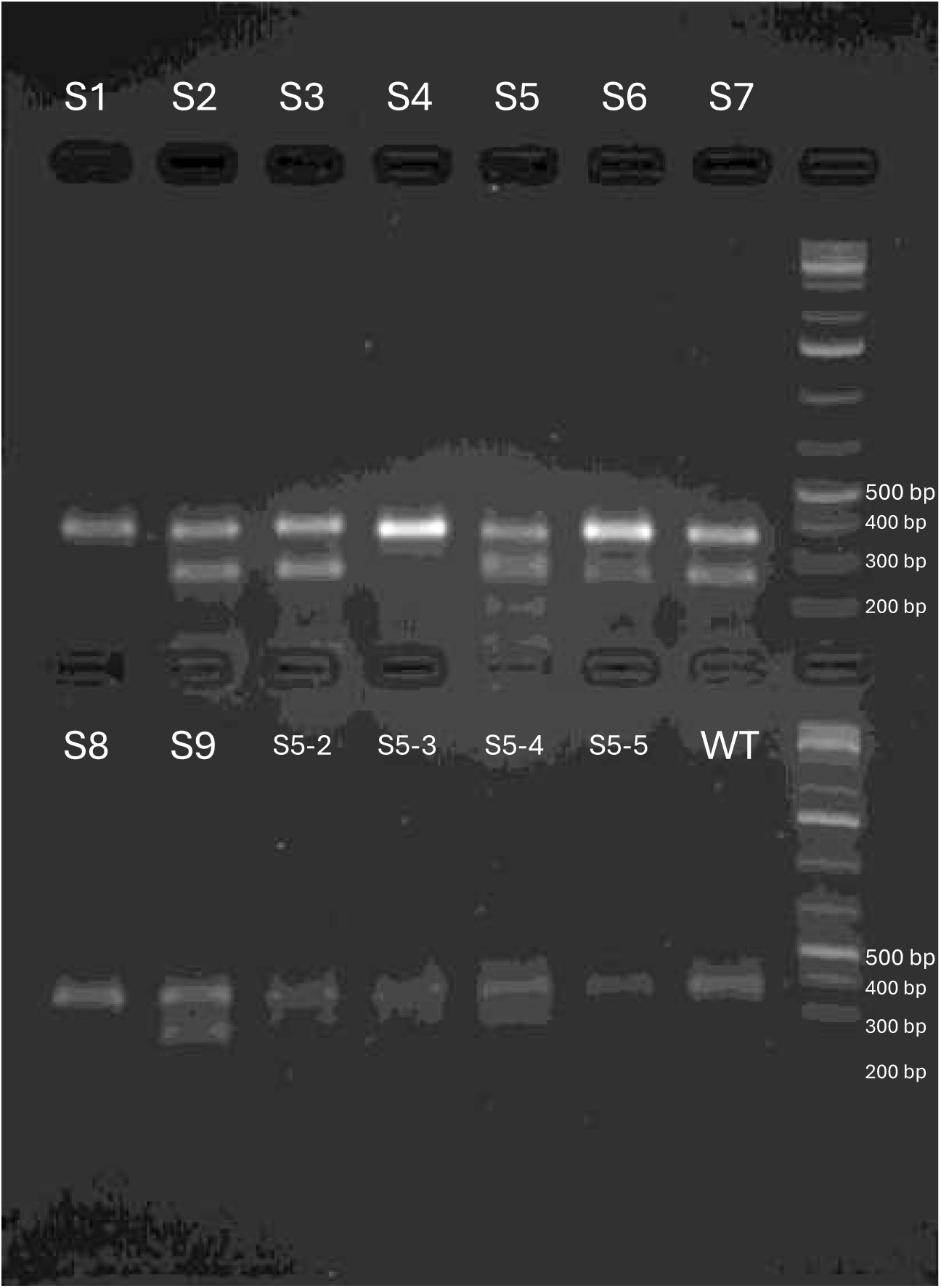
T7 endonuclease assay on *DNMT3B* mutant screening. S2, S3, S5, S6, S7, S9 are positive for mutation. S5-2 to S5-5 are the first brood offspring of S5. S1-S9 are pooled Brood 1 from their respective parent.

**Table 5.**
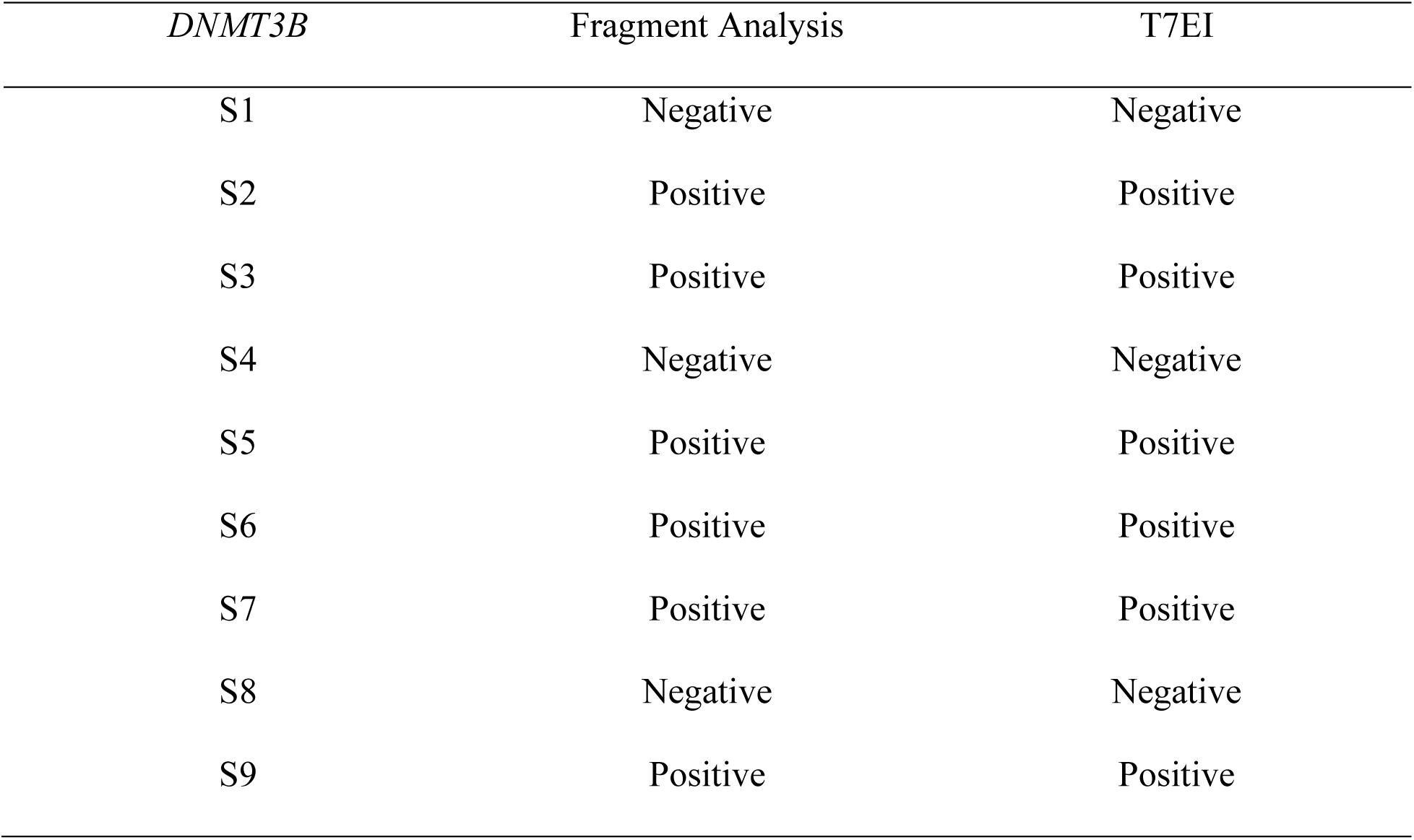
Mutant screening of S1-S9 for *DNMT3B* gene using Fragment Analysis and T7 endonuclease assay. Positive refers to a mutation detected and negative as no mutation.

| *DNMT3B* | Fragment Analysis | T7EI |
| --- | --- | --- |
| S1 | Negative | Negative |
| S2 | Positive | Positive |
| S3 | Positive | Positive |
| S4 | Negative | Negative |
| S5 | Positive | Positive |
| S6 | Positive | Positive |
| S7 | Positive | Positive |
| S8 | Negative | Negative |
| S9 | Positive | Positive |

### Case study #4 – *DMRT1* gene knockout

A total of 10 pooled samples, each representing a first brood offspring experiments knocking out the *DMRT1* gene, were screened for mutations. Fragment analysis detected mutations in 3 samples (Table 6). In contrast, T7EI cleavage assays showed positive results for all 10 samples, suggesting that all harbored some form of mutation (Figure 4).

**Figure 4.**
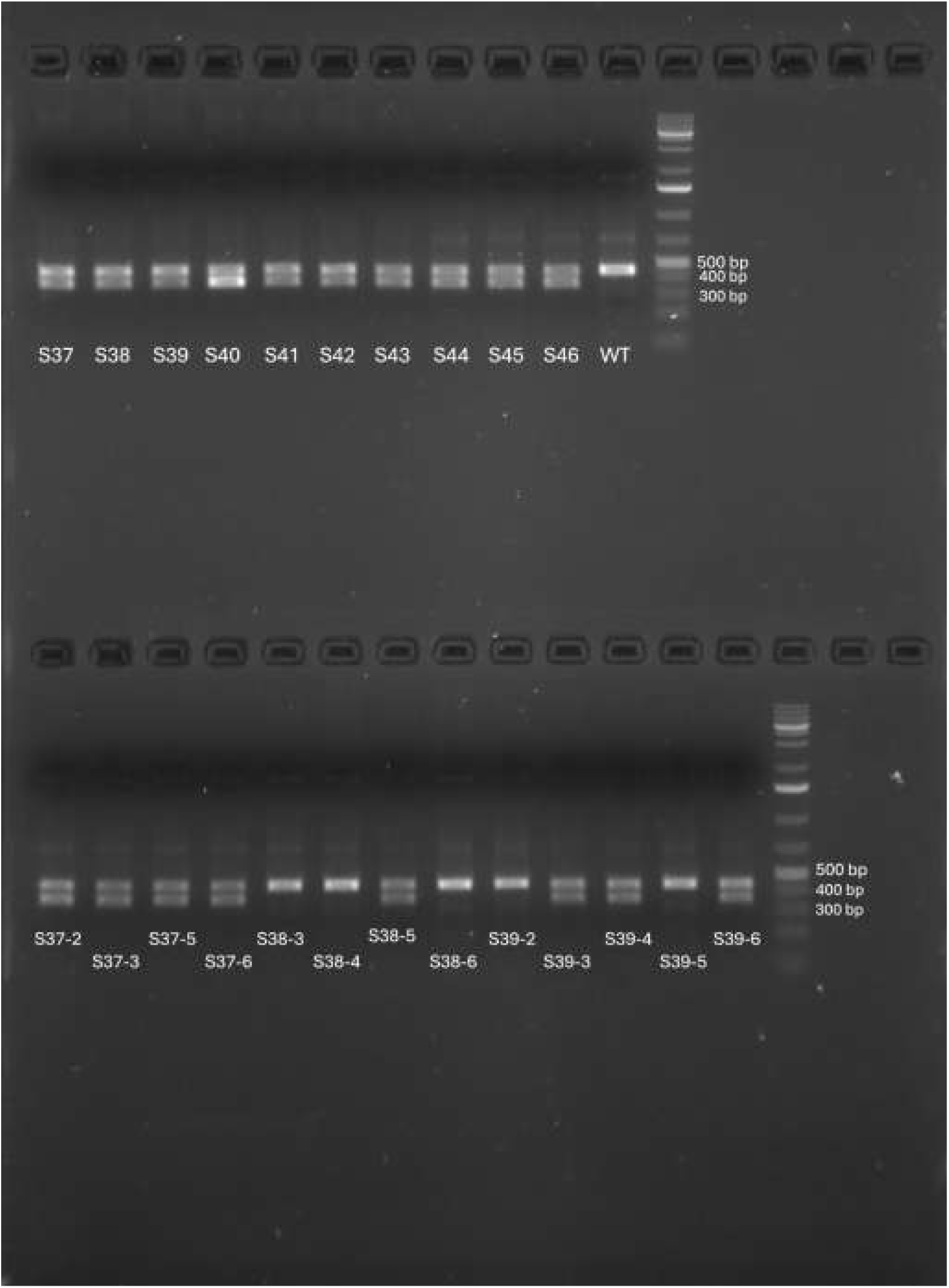
T7 on pooled DMRT1 gene for samples S37-S46, all are positive for T7 assay. T7 assay on individual brood 1 genotype from S37, 38 and S39. S38-3, S38-4, S38-6, and S39-5 showed no mutations, which is consistent with FA analysis result. T7 assay indicated no mutation for s39-2, conflicting the FA analysis result.

**Table 6.**
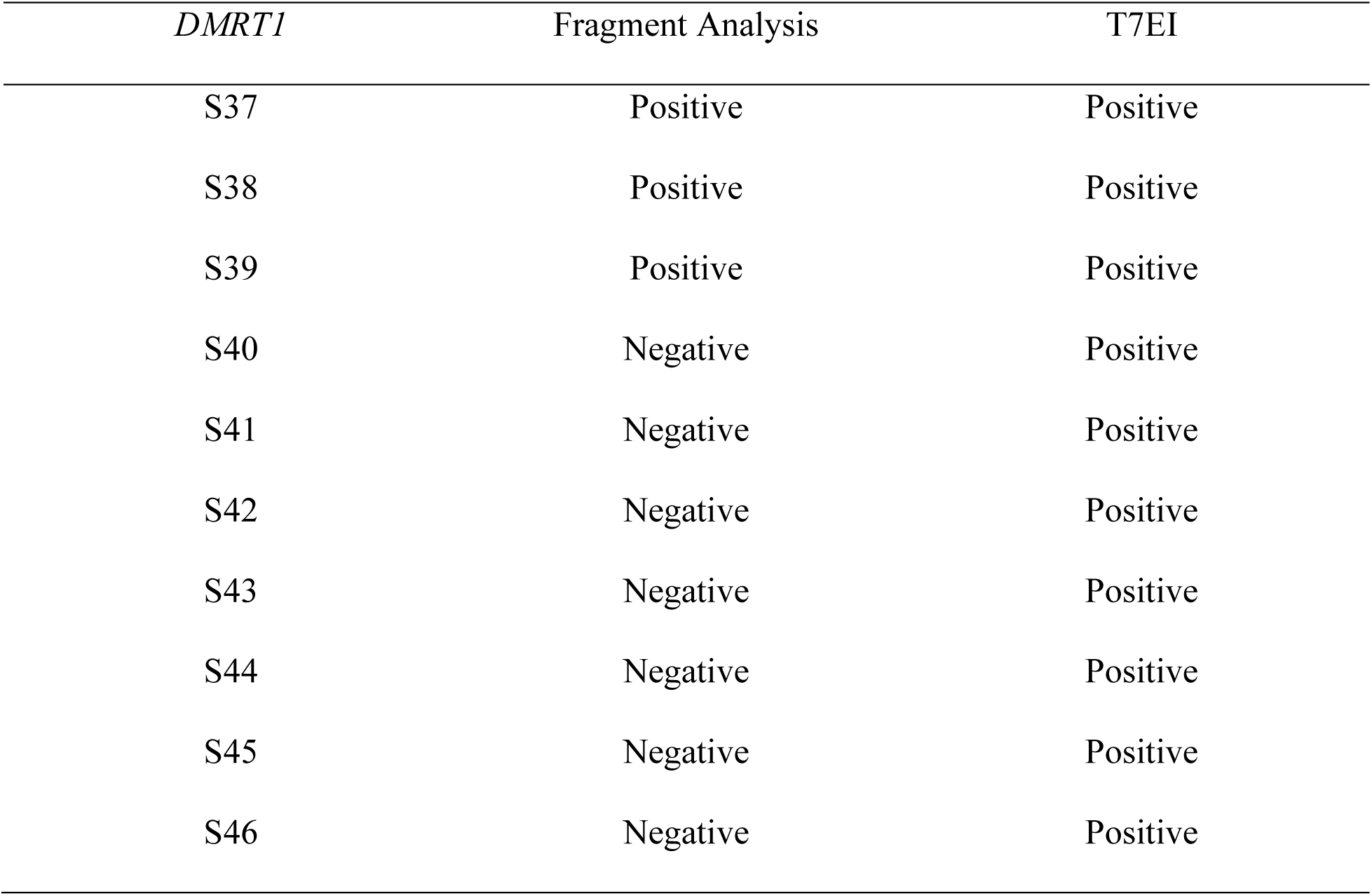
Mutant screening of S37-S36 for *DMRT1* gene using Fragment Analysis and T7 endonuclease assay. Positive refers to a mutation detected and negative as no mutation.

| *DMRT1* | Fragment Analysis | T7EI |
| --- | --- | --- |
| S37 | Positive | Positive |
| S38 | Positive | Positive |
| S39 | Positive | Positive |
| S40 | Negative | Positive |
| S41 | Negative | Positive |
| S42 | Negative | Positive |
| S43 | Negative | Positive |
| S44 | Negative | Positive |
| S45 | Negative | Positive |
| S46 | Negative | Positive |

**Table 7.** Mutant screening of individual Brood 1 from S37, S38 and S3 for *DMRT1* gene using Fragment Analysis and T7 endonuclease assay. Positive refers to a mutation detected and negative as no mutation.

| *DMRT1* | Fragment Analysis | T7EI |
| --- | --- | --- |
| S37-2 | Positive | Positive |
| S37-3 | Positive | Positive |
| S37-5 | Positive | Positive |
| S37-6 | Positive | Positive |
| S38-3 | Negative | Negative |
| S38-4 | Negative | Negative |
| S38-5 | Positive | Positive |
| S38-6 | Negative | Negative |
| S39-2 | Positive | Negative |
| S39-3 | Positive | Positive |
| S39-4 | Positive | Positive |
| S39-5 | Negative | Negative |
| S39-6 | Positive | Positive |

However, upon careful comparison with fragment analysis results, T7EI falsely identified 7 pooled samples as containing mutants. When T7EI was subsequently performed on individual (non-pooled) first brood samples, it was able to correctly show negative results for those that did not contain mutations (Figure 4), highlighting the limitation of T7 assays.

### Sensitivity and Accuracy

When comparing the two methods, FA demonstrated superior sensitivity in detecting small indels that were often missed by T7EI. The ability of capillary electrophoresis to resolve single-base pair differences provided an advantage over the enzymatic cleavage method, which relies on imperfect heteroduplex recognition. In contrast, T7EI was found to be more effective for screening large deletions but struggled with smaller mutations (see manual for Cat. No. M0302, New England Biolabs). T7 assay also shows different efficiency with different mismatches (Guan et al. 2004). This limitation makes FA the preferred method for applications requiring high-resolution indel analysis.

### Reproducibility and Practical Considerations

Reliability and reproducibility are critical for mutation detection. FA showed high reproducibility across multiple independent experiments, with minimal variation in fragment sizing and less optimization required. T7EI, however, exhibited inconsistent cleavage efficiency depending on sequence context, leading to variability in results. Additionally, the requirement for gel electrophoresis in T7EI introduces subjective interpretation, whereas FA provides objective numerical data for sizing of the alleles.

From a practical standpoint, T7EI remains an attractive option for laboratories with limited resources due to its lower cost and ease of implementation. It allows for rapid screening of large sample sets without the need for specialized instrumentation. On the other hand, FA, while requiring advanced equipment, provides more precise and reliable mutation characterization, making it an excellent choice for confirmatory analysis.

While sequencing remains the benchmark for mutation confirmation due to its ability to provide exact nucleotide changes (Knott and Doudna 2018), it is often impractical for handling large sample sets due to cost and time constraints. In such cases, FA and T7EI serve as feasible alternatives for high-throughput screening. FA, with its high resolution, offers a reliable method for detecting small indels, whereas T7EI provides a rapid and cost-effective option for initial screening although with higher chances of capturing false negatives. The choice between these methods depends on the specific needs of the study, with sequencing being the ideal validation tool when resources permit.

The T7 endonuclease 1 (T7E1) mismatch detection assay is widely utilized for assessing CRISPR-Cas9 editing efficiency. However, studies comparing T7E1 results with targeted next-generation sequencing (NGS) have demonstrated discrepancies, with T7E1 often failing to accurately reflect the true editing efficiency (Sentman et al.). Moreover, other alternative methods such as Tracking of Indels by Decomposition (TIDE) and Indel Detection by Amplicon Analysis (IDAA) have shown greater concordance with NGS data. These findings highlight the limitations of T7E1 in detecting certain mutation profiles and suggest that while useful for initial screening, it may not always provide reliable editing estimates.

## Conclusion

In this study, we compared two commonly used methods for mutant screening: the T7 endonuclease I assay and fragment analysis. Our findings highlight the strengths and limitations of each approach. T7EI is a cost-effective and rapid screening tool suitable but lacks sensitivity for detecting small mutations and is prone to false negatives. FA, while requiring specialized equipment, provides superior accuracy and resolution, making it the preferred choice for precise mutation analysis.

Ultimately, the choice of method depends on the specific requirements of a study. Laboratories seeking a quick and affordable screening method may benefit from T7EI, whereas those requiring high-resolution, higher sample quantities and a demand to quantitative mutation detection should consider fragment analysis. By understanding the trade-offs between these techniques, researchers can select the most appropriate tool for their CRISPR screening needs, particularly in whole-organism studies like our study conducted in *Daphnia*. We conclude that T7EI assay requires frequent optimizations based on amplicon length and composition, and types of mutation to detect (single base mismatches to large indels, heterozygous and homozygous mutations). In contrast, fragment analysis needs very little optimization except for optimizing the annealing temperature in PCR to reduce spurious amplifications. Fragment analysis also provides single base pair resolution when it comes to identifying mutant alleles.

## Data Availability Statement

The authors affirm that all data necessary for confirming the conclusions of the article are present within the article, figures, and tables.

## Acknowledgements

This work is supported by NSF EDGE program grants 2324639 (SX) and 2220696 (MEP), an NSF Career program grant 2348390 (SX), and a NIH grant R35GM133730 (SX). NIH grant R35GM133730 (SX)

